# Genomic and phenotypic studies among *Clostridioides difficile* isolates show a high prevalence of clade 2 and great diversity in clinical isolates from Mexican adults and children with healthcare-associated diarrhea

**DOI:** 10.1101/2024.02.01.578430

**Authors:** D Meléndez Sánchez, Laura Hernández, Miguel Ares, A Méndez Tenorio, Lourdes Flores-Luna, Javier Torres, M Camorlinga-Ponce

## Abstract

*Clostridioides difficile* is the most common cause of healthcare-associated infection worldwide. Its pathogenesis is mainly due to the production of toxins A, B, and CDT, whose genetic variants may be associated with disease severity. We studied genetic diversity in *C. difficile* isolates from hospitalized adults and children using different gene and genome typing methods and investigated their association with in vitro expression of toxins.

Whole-genome was sequenced in 39 toxigenic *C. difficile* isolates and used for multilocus sequence typing (MLST), *tcdA*, and *tcdB* typing sequence type, and for phylogenetic analysis. Strains were grown in broth media, and expression of toxin genes was measured by real-time PCR and cytotoxicity was determined in culture assays.

We found that clustering after genome-wide phylogeny followed clade classification, although different clusters were identified within each clade. The toxin profile *tcdA+/tcdB+/cdtA+/cdtB*+ and clade 2/ST1 were the most prevalent among isolates from children and adults. Isolates presented two TcdA and three TcdB subtypes, of which A2 and B2 were dominant. Toxin gene expression or cytotoxicity were not associated with genotyping or toxin subtypes. In conclusion, genomic and phenotypic analysis shows high diversity among *C. difficile* isolates from patients with healthcare-associated-diarrhea.

**Importance:** *Clostridioides difficile* is a toxin-producing bacterial pathogen associated with healthcare, and different genetic or phenotypic typing have been proposed for classification. We extensively studied cytotoxicity, expression of toxins, whole genome phylogeny, and toxin typing in clinical *C. difficile* isolates. Most isolates presented a *tcdA+ tcdB+cdtA+ cdtB+* pattern, with high diversity in cytotoxicity, and clade 2/ST1 was the most prevalent. However, they all had the same TcdA2/TcdB2 toxin subtype. Advances in genomics and bioinformatics tools offer the opportunity to understand the virulence of *C. difficile* better and find markers with better clinical use.

## Introduction

*Clostridioides difficile* is currently considered one of the most important causes of healthcare-associated diarrhea, although the number of community-based cases is increasing (1). *C. difficile* is a Gram-positive, anaerobic, spore-forming bacillus widely distributed in the intestinal tract of humans, animals, and the environment (2). Shortly after the identification of hypervirulent *C. difficile* strains RT 027 (B1/NAP1/ST1) in Canada, there were reports of CDI outbreaks caused by this strain in the United States and Europe and then Asia, Australia, and Latin America (3). In Mexico *C. difficile* RT-027 strain has been confirmed in hospitalized patients (4, 5, 6).

The clinical outcome of C. *difficile* infection (CDI) ranges from mild diarrhea to life-threatening pseudomembranous colitis, where virulence is mainly driven by the action of toxin A (TcdA) and toxin B (TcdB). TcdA and TcdB are 308 and 270 kDa proteins, respectively, belonging to a large Clostridial toxin family that glycosylate Rho family GTPases in host cells, leading to the disruption of the actin cytoskeleton, cell death, and a strong inflammatory response (7, 8). The genes that code for TcdA (*tcdA*) and TcdB (*tcdB*) are located within a 19.6 kb chromosomal region called the pathogenicity locus (PaLoc) (9,10). PaLoc contains five genes, two of which code for toxin A and toxin B, and three additional genes that are involved in the regulation, production, and secretion of toxins (*tcdR, tcdC*, and *tcdE*) (9, 10). Approximately 20% of *C. difficile* strains produce a third toxin called *C. difficile* transferase (CDT) or binary toxin, which is an ADP-ribosyltransferase encoded in a locus defined as the CdtLoc (11). CDT consists of *cdtA* and *cdtB* genes encoding the two subunits of CDT, plus *cdtR*, which encodes a positive regulatory protein (12).

Studies carried out with strains mutant in *tcdA* and *tcdB* in different animal models, along with the isolation of strains that produce only TcdB, have helped define that TcdA may contribute to disease severity, although TcdB alone can induce the full spectrum of disease in both animals and humans (13, 14). *C. difficile* strains may encode for different patterns of toxins, coding for one, two, three, or none of the toxins, primarily due to recombination and their highly mobile genome (15). Sequence analysis has revealed the existence of TcdA and TcdB variants (13,16) that are associated with biological properties such as enzymatic activity, immunogenicity, and affinity for receptors, suggesting variants may also be related to the severity of the disease (13, 16,17). Studies have identified twelve TcdB subtypes/variants (13), and strains like *C. difficile* 630 may present different TcdB variants, TcdB2, TcdB3, and TcdB4 (13, 18, 19) and up to seven subtypes/variants for TcdA (13).

In recent years, significant advances have been made in our understanding of the pathogenicity of *C. difficile* because of the identification and molecular characterization of the major toxins TcdA and TcdB. However, few studies have focused on the association of the pattern of coding toxin genes and toxins subtypes with expression levels and cytotoxicity of *TcdA, TcdB*, and *CdtA*. The possible association of genotyping of the strains (MLST, clade, and STs and whole-genome phylotyping) with toxin expression or cytotoxicity have not been studied either (9, 13). In addition, there is limited information on the subtypes of toxins in clinical isolates from pediatric patients. The aim of our study was to analyze toxins gene profiles, expression of TcdA, TcdB, and Cdt toxins, and the genotyping of *C. difficile* strains isolated from adult and pediatric hospitalized patients. For genotyping, MLST, clades, ST, and whole-genome phylogeny were determined and studied for any association with coding patterns, levels of expression, and variants of the *tcdA* and *tcdB* toxin genes.

## Results

### Amplification of toxin genes

Table 1 summarizes the characteristics of the patients and toxin genes profiles of the *C. difficile* strains analyzed. Thirty-three of the 39 *C. difficile* isolates (84.6%) were positive for all three toxin genes (*tcdA+ tcdB+cdtA+ cdtB+*). Other toxin profiles were less frequent: two (5.1%) were *tcdA-tcdB+cdtA-cdtB*-, three (7.7%) were *tcdA+ tcdB+ cdtA-cdtB*- and one isolate from a child was *tcdA-tcdB-cdtA-cdtB+*. The toxin pattern *tcdA+ tcdB+cdtA+ cdtB+* was significantly more frequent in isolates from adults than in isolates from pediatric patients (p<0.05).

**Table 1.** Characteristics of *Clostridioides difficile* strains isolated from children and adults with health-care associated diarrhea in Mexico.

| **Characteristics** | Adults<br>n=31<br>no. (%) | Children<br>n=8<br>no. (%) | Total<br>N=39<br>no. (%) |
| --- | --- | --- | --- |
| **Age, mean ±SD** | 57.4±14.7 | 6.2±6.0 | 35.2±25.0 |
| **Gender** |  |  |  |
| Male | 14 (45.1) | 5 (62.5) | 19 (48.7) |
| Female | 17 (54.9) | 3 (37.5) | 20 (51.2) |
| **Toxin genes profile** |  |  |  |
| *tcdA+* *tcdB+* *cdtA+* *cdtB+* | 28 (93.5) | 5 (62.5) | 33 (84.6) |
| *tcdA-* *tcdB+* *cdtA-* *cdtB-* | 2 (6.5) | 0.0 | 2 (5.1) |
| *tcdA+* *tcdB+* *cdtA-* *cdtB-* | 1 (3.2) | 2 (25.0) | 3 (7.7) |
| *tcdA-* *tcdB-* *cdtA-* *cdtB+* | 0.0 | 1 (12.5) | 1 (2.5) |

### Production of toxins *in vitro*

Once we learned about the toxin gene profiles, we asked whether there was any difference in the cytotoxic activity of strains after in vitro culture. As shown in Figure 1A, the supernatant of the adult *C. difficile* strains clearly showed higher cytotoxicity titers than the isolates from children.

**Figure 1.**
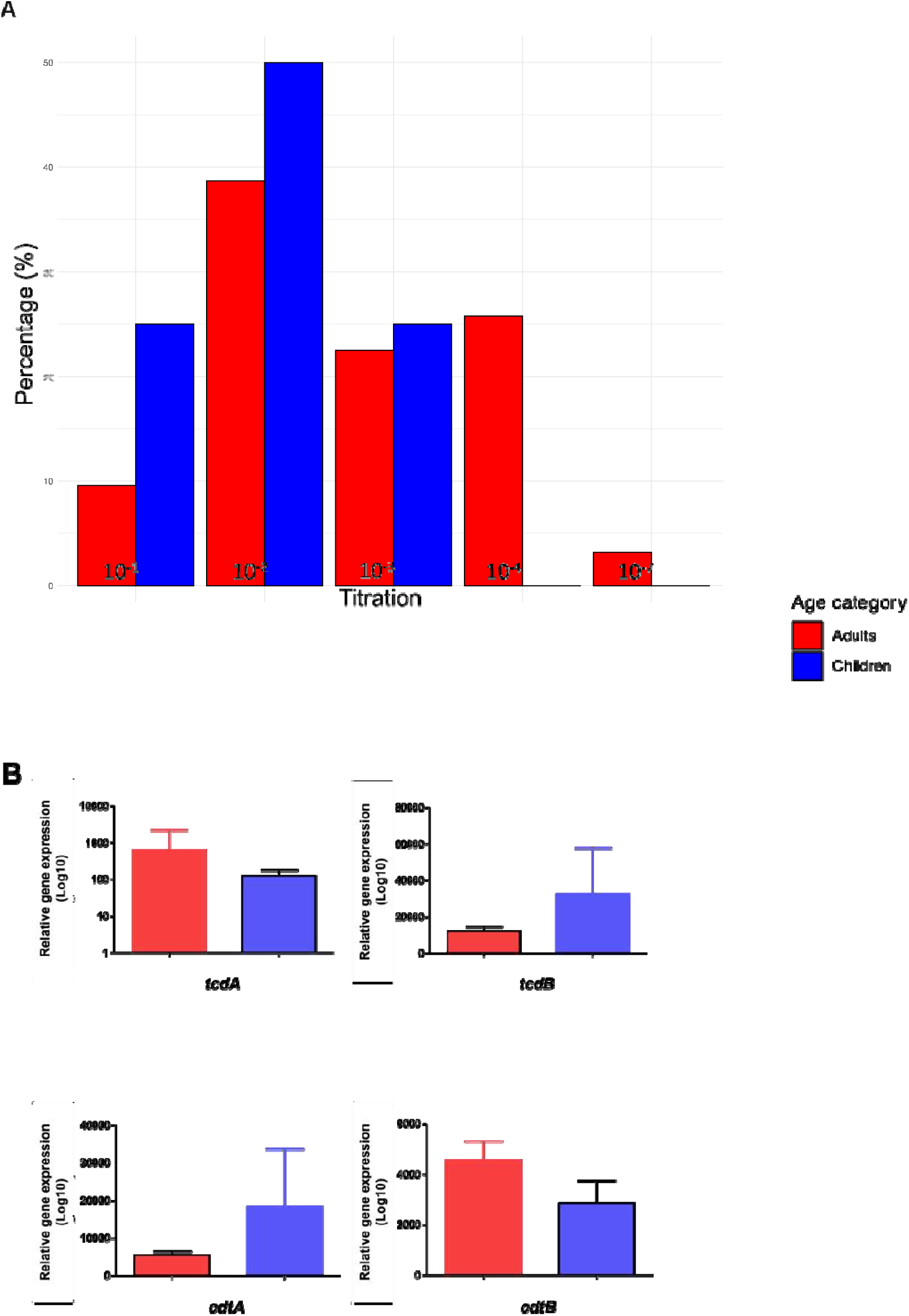
Titration of TcdB of *C. difficile* and toxin gene expression analysis for TcdA, TcdB, CdtA, and CdtB. Figure 1A.*C. difficile* toxin B titration on Vero cells after 24 h incubation Figure 1B.Toxin gene expression of each toxin (TcdA, TcdB, CdtA and CdtB). Means and standard deviation from triplicates are shown. Each toxin gene expression was normalized using *rpoA* as a reference.

#### Toxin Gene expression

Differences were observed in the relative expression of toxin genes by strains isolated from adults vs. children (Figure 1B); whereas adults expressed more *tcdA* than children, children expressed more *tcdB* than adults (Figure 1B). Similarly, whereas *cdtA* was expressed more in children, *cdtB* was more expressed in adults (Figure 1B). Thus, the relative expression *tcdA/tcdB* was 4.9 in adults and 0.37 in children’s strains. In contrast, the relative expression of *cdtA/cdtB* was 0.29 in adults and 1.29 in children’s strains.

#### Genotyping analysis

Moving to the genotyping of the 39 *C. difficile* isolates, we first analysed of MLST data, which showed the presence of clades 1, 2, and 4, including ten different sequence types (STs) (Table 2). Thirty-three (85%) isolates were clade 2; among them 29 were ST1 (74% of the 39 strains were clade 2/ST1). All other ST types were present in a single strain, except for ST37, present in two adult isolates with the pattern (*tcdA-tcdB+ cdtA-cdtB-)* that belonged to clade 4, whereas the single strain (*tcdA-tcdB-cdtA-cdtB+*) was isolated from a child and was clade 2.

**Table 2.** Distribution of toxin profile, clades, and sequence type, in 39 *Clostridioides difficile* strains isolated from adults and children in Mexico.

| Toxins profile | Clade | Sequence type (ST) | Adults<br>n=31 | Children<br>n=8 |
| --- | --- | --- | --- | --- |
| *tcdA+* *tcdB+*<br>*cdtA+* *cdtB+* |  |  | No (%) |  |
|  | 2 | 1 | 24(77.4) | 5(62.5) |
|  | 1 | 2 | 1(3.2) | 0.0 |
|  | 1 | 42 | 1(3.2) | 0.0 |
|  | 2 | 48 | 1(3.2) | 0.0 |
|  | 2 | 95 | 1(3.2) | 0.0 |
| *tcdA-* *tcdB+*<br>*cdtA-* *cdtB-* | 4 | 37 | 2(6.4) | 0.0 |
| *tcdA+* *tcdB+*<br>*cdtA-* *cdtB-* | 1 | 35 | 0.0 | 1(12.5) |
|  | 2 | 41 | 0.0 | 1(12.5) |
|  | 1 | 55 | 1(3.2) | 0.0 |
| *tcdA-* *tcdB-*<br>*cdtA-* *cdtB+* | 2 | 708 | 0.0 | 1(12.5) |

#### Subtyping of TcdA and TcdB

The 39-genome sequences from *C. difficile* clinical isolates were analyzed for TcdA and TcdB toxin subtypes. We found two TcdA subtypes and three TcdB subtypes in the Mexican isolates (Table 3). In the analyses of TcdA most of the strains (71.4%) had an A2.1 subtype. Analysis of TcdB subtypes showed that B2.1 was the predominant subtype (76.3%), although other subtypes were also identified. The subtype A1.1/B1.2 was the dominant toxins subtype. Also, we examined the correlation between the TcdA and TcdB subtypes and the *C. difficile* whole-genome phylogeny and observed a strong association between the toxins subtype and clades. All clade 2 were Tcd A2/B2, the four clade 1 were Tcd A1/B1, and the two clade 4 were Tcd A1/B3 (Figure 2B).

**Table 3.** Distribution of TcdA and TcdB subtypes of Mexican *C. difficile* strains.

| Subtype A | Strains<br>No(%) | Subtype B | Strains<br>No(%) |
| --- | --- | --- | --- |
| A1.1* | 3 (8.5) | B1.2** | 2 (5.2) |
| A1.8 | 1 (2.8) | B1.3 | 1 (2.6) |
| A2.1 | 25 (71.4) | B1.4 | 1 (2.6) |
| A2.14 | 3 (8.5) | B2.1 | 29 (76.3) |
| A2.3 | 1 (2.8) | B2.19 | 2 (5.2) |
| A2.4 | 2 (5.7) | B2.22 | 1 (2.6) |
|  |  | B3.1*** | 2 (5.4) |
| Total | 35 | Total | 38 |

\* Test of proportions, $p < 0.01$ the proportion different between the sum of A1.1, A1.8 and the sum of A2.1, A2.14, A2.3, A2.4 .
\*\* Test of proportions, $p < 0.01$ the proportion different between the sum of B1.2, B1.3, B1.4 and the B2.1, B2.19, B2.22.
\*\*\*Test of proportions, $p < 0.01$ the proportion different between the sum of B2.1, B2.19, B2.22 and B3.1.

**Figure 2.**
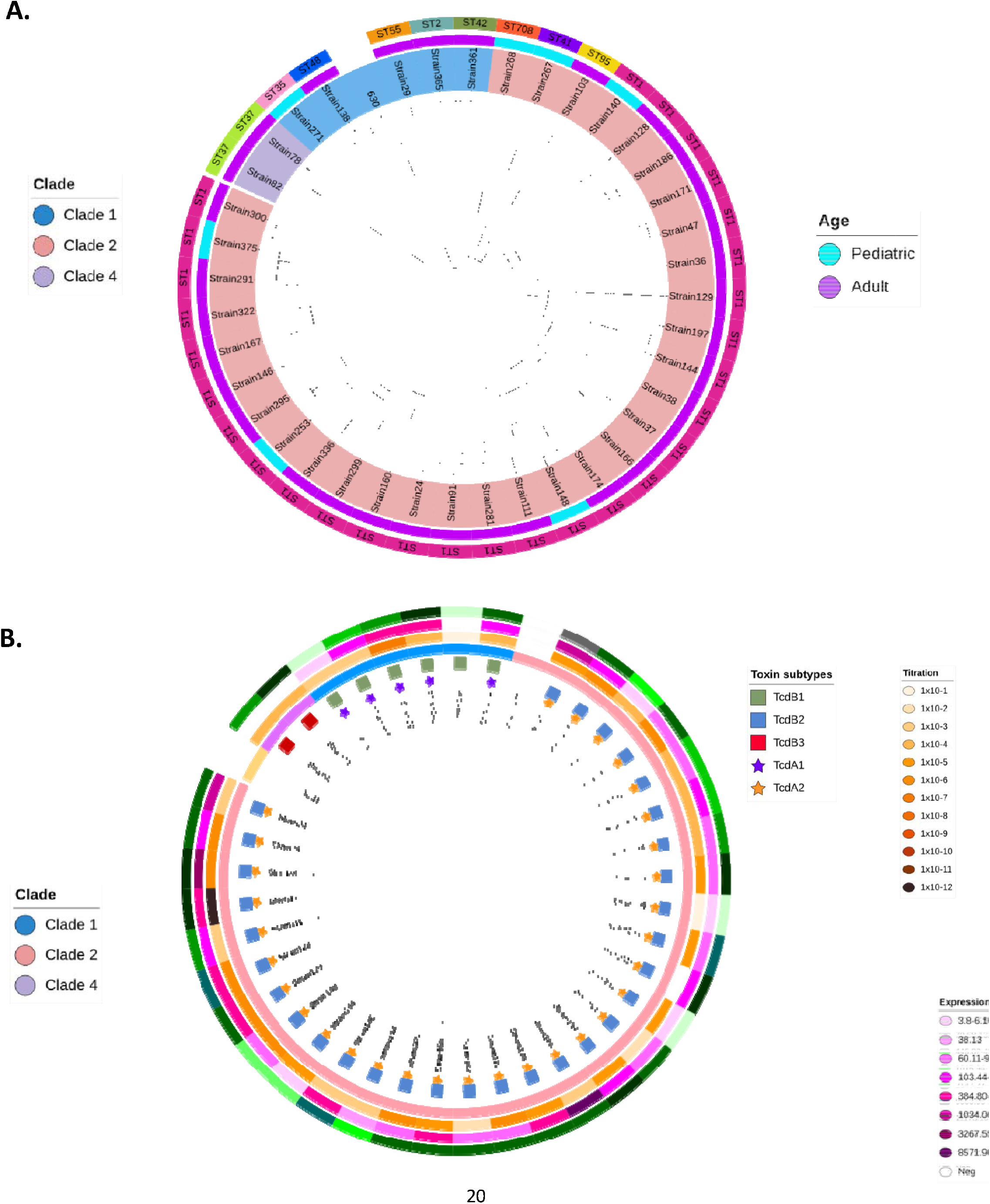
Genome sequence analysis of 39 *C. difficile* isolates, their clade distribution, sequence type(ST), and TcdA and TcdB toxin subtypes. Figure 2A. Complete genome sequence analysis of 39 *C. difficile* isolates from 31 adults and 8 children. The Phylogenetic Tree was created with MLST analysis, with the Vamphyre software; the STs are in the outer ring marked with different colors. The clades are represented by three different colors (C1 blue, C2 pink, and C4 light purple). The age of the patient is indicated in the second ring by two different colors (adult purple and children’s blue). *C. difficile* 630 was used as a control. Figure 2B. Phylogenetic analysis of the TcdA and TcdB subtypes of 39 *C. difficile* isolates from Mexican patients. The tree was built with FastTree V2.1.11 using the maximum likelihood distances according to the Jones-Taylor-Thorton evolution model and visualized with iTOL v.6.7.5 (https://itol.embl.de/). Marked at nodes withstar symbols for TcdA subtypes and boxes for TcdB subtypes. The next ring shows the clade to which each studied sequence belongs. The result of the TcdB titration is shown in the following ring marked in orange and brown tones; 291* strain did not undergo TcdB titration. The expression values of TcdA are observed in the next ring marked in purple and in the outermost ring, the expression of TcdB marked in green tones is observed. Strain 268*** had the toxin pattern: *tcdA-tcdB-cdtA+*. As an external group, the hemorrhagic toxin TcsH from *C. sordelli* was used.

### Genome phylogenetic analysis

Genome phylogenetic analysis was performed with the whole-genome sequence of the 39 *C. difficile* strains. The tree split into 3 main branches, each with strains belonging to a unique clade, 2, 1, or 4 (Figure 2A). However, several clusters were formed within each main branch, mainly for clade 2, where many subclusters were observed, even within the 31 clade 2/ST1 strains, evidencing large genomic distances within them. In some instances, unusual ST types were associated with subclusters, like ST708, ST41, and ST95 within the large clade 2 group. In the few strains within the clade 1 group, subclusters included different ST types. The whole-genome analyses did not distinguish between isolates from adults and children, although unusual ST types were more common in strains from children (Figure 2A).

### Phylogenetic analyses with concatenated TcdA and TcdB amino acid sequences

We next asked if there was any association between the phylogeny of TcdA and TcdB genotypes with clades, ST, cytotoxicity titers, or gene expression (illustrated in Figure 2B). The concatenated amino acid sequences of TcdA and TcdB were used to construct a phylogenetic tree, using *C. sordellii* TcsH toxins as an external reference. Phylogeny of TcdA and TcdB showed a clear clustering according to clades; thus, all clade 2 strains were TcdA2/TcdB2, except one nontoxigenic (strain 268). Also, all clade 1 strains were TcdA1/TcdB1, and those clade 4 were TcdA1/TcdB3. Still, even within each clade and toxin genotype, there was sequence diversity in the toxin genes as evidenced in the tree (Figure 2B). Thus, whereas 25 of the 31 clade 2-TcdA2/TcdB2 strains fell into a subcluster with almost no distance between them (Figure 2B, from Strain 24 to Strain 336) the other six were in separated clusters, three were ST1 type whereas the other three showed uncommon ST’
ss (08, 41 and 95). It is of interest to note that three of these six strains were isolated from children. In addition, in clades 4 and 1, the toxins sequence was diverse among all strains. Thus, whereas clades are strongly correlated with toxin types, sequences of TcdA and TcdB present diversity.

No correlation of toxins genotype with cytotoxicity or toxin gene expression was observed (Figure 2B). There was a high diversity in cytotoxicity and expression of genes, even within the 28 strains clade 2-ST1-TcdA2/TcdB2, with a range from Strain 299 with 10^−1^ cytotoxicity/<6.0 *tcdA* expression/<10 *tcdB* expression to Strain 47 with 10^−12^ cytotoxicity/500 *tcdA* expression/>20,000 *tcdB* expression. Also, cytotoxicity and toxin expression were variable among the six clade 1/TcdA1/TcdB1 strains.

## Discussion

During the previous *C. difficile* pandemic in the 80s-90s we learned on the main role of TcdA and TcdB in the pathogenesis of the infection, but little was known about the molecular epidemiology of the infection. At that time, strains coding for only one toxin (*tcdA-/tcdB*+) were very rare (20). In the past 20 years, a second pandemic extended worldwide, and thanks to updated genomic techniques, we are now learning more in detail on the molecular epidemiology of the infection. *C. difficile* clade 2/ST1 strain has spread throughout the world, including Latin America (1), although in recent years, the incidence of CDI, particularly by clade 2/ST1, seems to have decreased (21,22). However, the epidemic keeps evolving, and new clades have emerged with a wide diversity of genotypes in different countries (Clades 1, 2, 3, 4, and 5 and one *cryptic* clade have been described) (23,24). Studies in large numbers of strains have shown that clade 1 is more prevalent worldwide, whereas, in Mexico, we found clade 2/ST1 strains to the more prevalent, in isolates from both adults and children (6). Similarly, Rohana et al. (25) reported that clade 1, ST04, and ST037 strains were the most frequent in Israel. It is interesting to note that we found clade 4 and clade 5 strains with unusual STs (37, 35, 41, 48, 55, 95, and ST708) among Mexican isolates. Some of these STs have been isolated from different environmental, animal, and clinical sources (26) but are now circulating among humans and emerging as clinically important strains.

*C. difficile* isolates of the molecular type Clade2/ST1 have been associated with severe disease and in hospital outbreaks, and it has been suggested that these “hypervirulent” strains produce higher levels of TcdA and TcdB toxins besides the CDT toxin (28).In the present work, we found clade 2/ST1 as the most prevalent and extensively studied cytotoxicity, toxins profile, toxins expression, and whole-genome phylogeny.As suggested, most of these clade 2/ST1 strains presented the toxins profile *tcdA+ tcdB+cdtA+ cdtB+*. Still, they showed a high diversity in cytotoxicity and expression of both *tcdA* and *tcdB*, although they all had the same toxins subtype TcdA2/TcdB2.In agreement with our findings, recent evidence suggests that clade 2/ST1 isolates do not necessarily produce more toxins *in vitro* than isolates from other lineages (28), as was observed in early reports with hypervirulent *C. difficile* 027 (29). Watanabe et al. also found no association between clades and toxin gene expression (27).

Evidence of extreme homologous recombination in the *tcdB* gene has been found, suggesting a strong positive selective pressure, resulting in diversification of the gene sequence (16). These findings are consistent with our results and may explain the greater number of subtypes that we identified in the analyzed sequences (Table 3), 3 subtypes in TcdB compared to 2 subtypes in TcdA. It was interesting to observe that the whole-genome phylogenic analyses clearly grouped clade 2/ST1 strains into different clusters (figure 2A), suggesting the presence of differences in other regions of the genome that may also influence virulence, like outer membrane proteins or sporulation genes (12).A deeper analysis of the genomes may identify genes or regions responsible for differentiating the clades, but also to find the simplest and most efficient test for molecular epidemiology studies and to better identify the virulence of the strains.

This report also provides insight into the genotyping and expression of toxins in isolates from children and adults, whose possible differences are poorly described. Although the number of children strains was low, some differences were observed. The expression levels of *tcdA, tcdB, cdtA*, and *cdtB* were markedly different compared to adults isolates (Figure 1). In addition, the presence of rare STs was common in children isolates (ST41, 708, and 35) (do children often acquire *C. difficile* from the environment?), and the only strain clade 2 that had no *cdtA* and *cdtB* genes was from a child (Strain 268, clade 2/ST708).

Our analysis of toxin subtypes identified three subtypes for Tad and two subtypes for TcdA as prevalent in our community, but additional subtypes will most probably be identified when studying a more significant number of strains, as illustrated by Shen *et al*. examining strains from different geographic regions (13). TcdB2 was our patients’ most frequently identified subtype, unlike studies in other geographical areas where TcdB1 was the most common subtype (13, 16). The study of toxin subtype is relevant; whereas initial studies reported that TcdB1 strains might be more cytotoxic, more recent analyses have suggested that TcdB2 was more cytotoxic in other populations (19). Furthermore, it has been found that sequence variations between subtypes may have a negative impact on therapeutic treatment with antibodies and vaccines (16). Clearly, more studies on toxins typing and virulence are needed. We observed that, for example, not all tcdA2/tcdB2 are homologous, and different clusters are observed when analyzing genetic distance (Figure 2) that may have implications on the pathogenic activity.

A clear limitation of this study is the low number of isolates tested, which limit the reach of our findings, particularly in strains isolated from children. Still, the genomic and phenotypic results of our work offer clues on the diversity of *C. difficile* strains circulating in one community. Also, no clear correlation was found between genotypic and phenotypic properties, e.g., the cytotoxicity and expression of toxin genes varied even within clade 2/ST1 strains, and no evidence of uniform high toxicity was revealed, as previously suggested. During the first pandemic, we used to classify strains as toxigenic or non-toxigenic and thought that *C. difficile* was more of a clonal species. In this second pandemic, advances in genomics and bioinformatics tools offer the opportunity to go further and extend our studies to all genome content to understand virulence better and find markers with better clinical use.

## Material and Methods

### Patients and isolation of *Clostridiodes difficile*

A total of 39 *C. difficile* strains isolated from patients with nosocomial diarrhea were analyzed, 31 isolated from adults and 8 from children. Patients were attended at the Centro Médico Nacional Siglo XXI, Instituto Mexicano del Seguro Social in Mexico City from 2016 to 2018. The ethical committee from the Instituto Mexicano del Seguro Social approved the study and patients or their guardians were informed about the study and asked to sign an informed consent for participation.

To isolate *C. difficile*, stool samples were treated with 96% ethanol at room temperature for 50 min, followed by centrifugation at 4000 rpm for 10 min (6). The pellets were inoculated onto taurocholate-cycloserine-cefoxitin-fructose agar (CCFA) and incubated at 37°C for 48h under anaerobic conditions in a jar with an atmosphere containing 85% N2, 5% H2, and 10% CO2 generated using an Anoxomat system (MART Microbiology BV; Netherlands). *C. difficile* colonies were identified by colonial morphology, Gram stain, fluorescence under UV light, odor, and Vitek MS, and confirmed by the amplification of *tpi* genes by PCR, using primers and amplification procedures previously described (30). All isolates were frozen at -70°C in Brucella broth medium supplemented with 10% glycerol for subsequent analysis.

### RNA extraction for qRT-PCR

*C. difficile* strains were grown in 5 mL of TY broth for 24 hs under an anaerobic atmosphere in a jar with an atmosphere containing 85% N2, 5% H2, and 10% CO2 generated using an Anoxomat system (MART Microbiology BV; Netherlands) at 37°C for 48h and RNA extracted from the bacterial pellet using the hot phenol method (31). Samples were chilled on ice, centrifuged at 19,000 × g for 10 min at 4 °C. The aqueous layer was transferred to a 1.5 ml Eppendorf tube, and RNA was precipitated with cold ethanol and incubated at ™70°C overnight. The RNA was centrifuged at 19,000 × g for 10 min at 4°C. Pellets were washed with cold 70% ethanol and centrifuged at 19,000 × g for 5 min at 4°C and air dried for 15 min in the Centrifugal Vacuum Concentrator 5301 (Eppendorf, USA). The quality of RNA was assessed using a NanoDrop (ND-1000; Thermo Scientific, USA). cDNA was synthesized using 1 µg of RNA, 0.2µg/µl of random hexamer primers, and 2 U/µl of M-MulV-RT (Reverse transcriptase of Moloney Murine Leukemia Virus; Thermo Scientific, USA).

### Expression of toxin genes measured by qRT-PCR

The expression of toxins A, B, and CDT genes was determined by quantitative reverse transcriptase PCR (qRT-PCR) in a LightCycler 480 thermal cycler (Roche Diagnostics, USA). The primers specific for toxin A, toxin B, and toxin CDT genes used are listed in Table S1(32, 33). Master Mix was prepared forward and reverse primers (20 µM), cDNA (50-100 ng), and SYBR Green Master 2x. A 96-multiwell plate containing all samples was loaded into the lightCycler. Each sample was tested in triplicate, and the reported relative expression represents the mean of the three replicates. The expression value of all toxin genes was normalized with the expression of the *C. difficile* housekeeping gene *rpoA* and used to estimate relative expression as follows: relative expression *=* 2^(CTrpoA-CT target gene)^. We selected *C. difficile* 630 and *C. difficile* 027 as control strains (16).

### In vitro *toxin production and cytotoxin titration assay*

*C. difficile* strains were cultured in Trypticase Yeast Extract broth (TY) and incubated at 37°C for 24-48 h in a jar under an anaerobic atmosphere. Culture supernatants were collected by centrifugation. Green monkey kidney fibroblast cells (Vero) were grown in 25 cm^2^ flasks with MEM media supplemented with 10% fetal serum (Gibco, USA) and incubated at 37°C in a 5% CO_2_ atmosphere. Aliquots of 200 ul with 5 × 10^3^ cells /mL were distributed per well in 96-well microtiter plates. Serial ten-fold dilutions of the test culture-supernatant were applied (10 ul) to confluent cells as previously described (6) and incubated for 24 hs at 37°C in a 5% CO_2_ atmosphere. Cytotoxic units (CU) were expressed as the inverse of the maximum dilution that caused rounding of at least 50% of the cells. For neutralization assays, samples were mixed with an equal volume of antiserum to toxin B (Techlab, USA) and incubated for 30 minutes before inoculation onto cells. Each assay was performed by duplicate.

### Whole–genome sequencing (WGS) and bioinformatics analysis

*C. difficile* strains were grown overnight in BHI broth, centrifuged, and genomic DNA extracted using the DNeasy Kit (Qiagen, Hilden, Germany). Genome was sequenced using a Hi Seq 2000 (Illumina Inc., San Diego, CA, United States) as described (6). Sequences were assembled *de novo* with default settings with the Shovill pipeline v (https://github.com/tseemann/shovill) (34). The contings were annotated using the annotation pipeline Prokka **v**.1.13(https://github.com/tseemann/prokka) (35). The 39 *C. difficile* genomes were previously sequenced in an earlier study (6). Genome sequences were deposited at the NCBI as part of the 100K Pathogen Genome Project (34) under the BioProject accession number PRJNA203445. Supplementary Table 1 describes the accession number for each genome sequence.

### Phylogenetic analyses of the whole genome sequence

The whole-genome sequence of the 39 *C. difficile* strains was performed using the Newick format file obtained with VAMPhyRE software (http://biomedbiotec.encb.ipn.mx/VAMPhyRE/), and the tree was edited with iTol (https://itol.embl.de/). The sequence of C. *difficile* ATCC 630 (GenBank AM180355.1) was used as a reference strain.

### Subtyping of TcdA and TcdB

For the identification of toxin variants, the *tcdA* and *tcdB* gene sequences were extracted from pre-computed genome annotations and extracted using probabilistic profile hidden Markov models (profile HMMs) with software HMMER V.3.3.2 (http://hmmer.org). The sequences were aligned with ClustalQ in Seaview v.4, using strain 630 as queries. Sequences were translated from nucleotides to amino acid sequences with Seaview v.4, and the subtype of toxins was determined with DiffBase database (http://diffbase.uwaterloo.ca)(19). The amino acid sequences of *tcdA* and *tcdB* were concatenated, and a phylogenetic tree was built with FastTree V2.1.11 using the maximum likelihood distances according to the Jones-Taylor-Thorton evolution model and visualized with iTOL v.6.7.5 (https://itol.embl.de/). The *C. sordelli* tcsH sequences were used as foreign genes for the analysis (16).

### Statistical analyses

A descriptive analysis was done, absolute and relative frequencies were obtained for categorical and the mean for continuous variables. The proportion test was used to evaluate the differences in the frequency of detection of toxin genes and sequence type between adults and children strains. The Wilcoxon rank-sum test evaluated differences in toxin gene expression between adults and children strains.

---

**Table 1 Supplementary.**
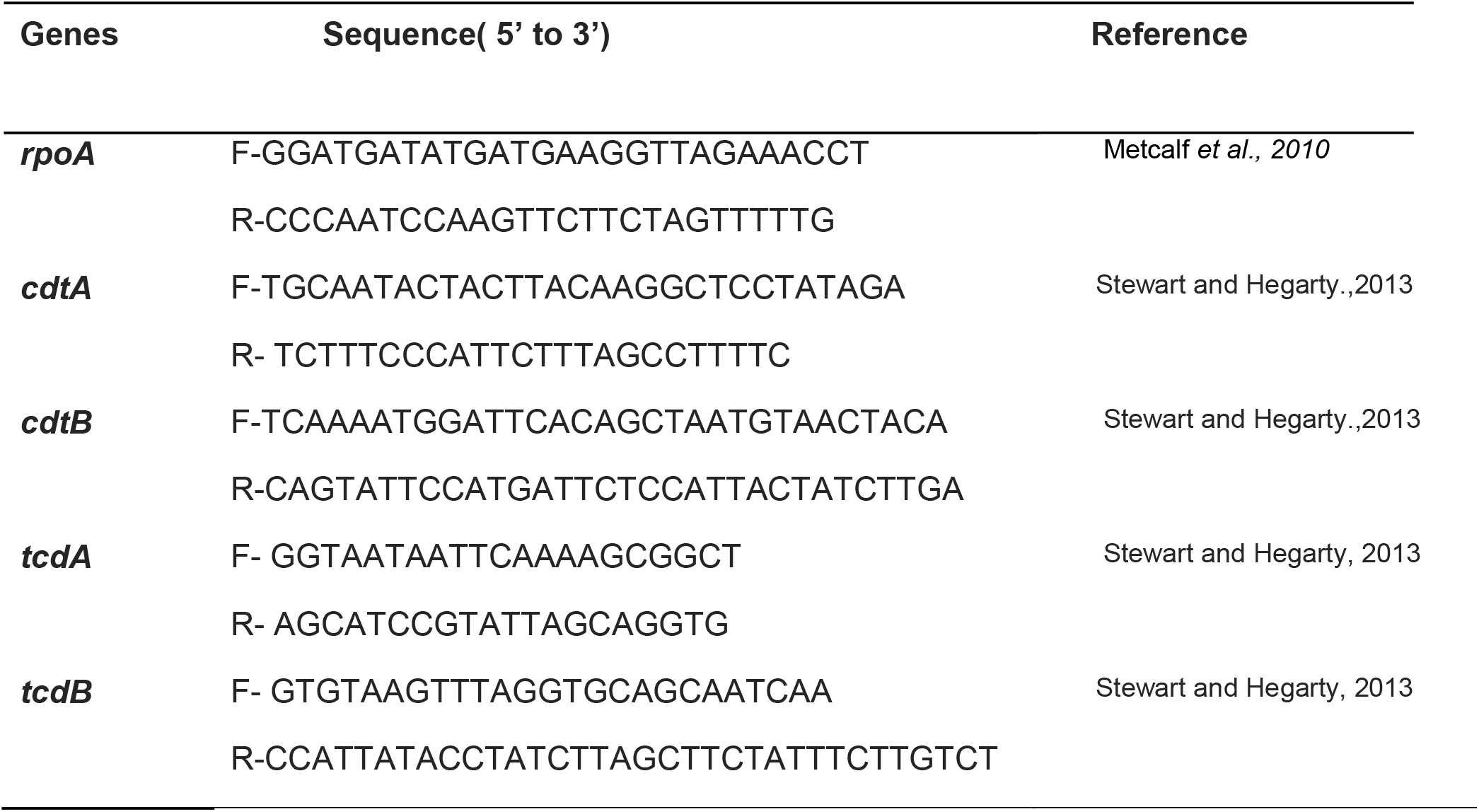
Primers used in this study.

